# Dynamic instability of force-generating bacterial microtubules

**DOI:** 10.1101/2023.08.02.551647

**Authors:** Reza Amini H., Vladimir A. Volkov, Marileen Dogterom

## Abstract

Dynamic instability refers to the ability of cytoskeletal polymers to switch between growing and shrinking phases. This phenomenon has been extensively studied for eukaryotic microtubules which consist of 13 protofilaments. Here we report on the dynamic properties of prokaryotic microtubules found in *Prosthecobacter* bacteria, which consist of 4-5 protofilaments and, like their eukaryotic counterparts, display dynamic instability. Using microfabricated barriers we show that the catastrophe rate of bacterial microtubules increases when their growth is stalled by a rigid barrier. We find that the lifetime distributions of both free and stalled bacterial microtubules can be fitted using the same phenomenological model that we previously introduced for eukaryotic microtubules, suggesting that bacterial microtubules may be considered a model system for eukaryotic microtubules. We further use cryo-electron tomography to reveal structural details of dynamic ends and show that bacterial microtubules may form doublets similar to axonemal microtubules in eukaryotes.

## Introduction

In eukaryotic cells, microtubules (MTs) are hollow cylindrical polymers that play essential roles in various cellular activities including cell division, cell motility and intracellular transport (reviewed in (Akhmanova & Steinmetz, 2008; Gudimchuk & McIntosh, 2021)). Microtubules alternate between growing and shrinking states *in vitro* and *in vivo*, a phenomenon known as dynamic instability (Cassimeris et al., 1988; Mitchison & Kirschner, 1984). Microtubules grow by the addition of GTP-bound tubulin to their ends. Subsequential hydrolysis of tubulin-bound GTP leads to a microtubule lattice that is primarily composed of GDP-bound tubulin, a situation that is inherently unstable. A growing microtubule may switch to rapid shrinking in an event termed a catastrophe when a stabilizing cap that is rich in unhydrolyzed GTP is lost from the microtubule end (Gudimchuk & McIntosh, 2021; Vleugel et al., 2016). The exact nature as well as the size of the stabilizing cap is still a topic of debate despite a long series of experimental and modeling efforts (Duellberg et al., 2016; Kok et al., 2021; McIntosh et al., 2018; Rickman et al., 2017; Roostalu et al., 2020; Zakharov et al., 2015).

Although dynamic instability is a hallmark of eukaryotic microtubules, prokaryotic tubulin-like filaments have recently been shown to display dynamic instability as well. Bacterial microtubules were for example discovered in some *Prosthecobacter* species (Deng et al., 2017; Jenkins et al., 2002; Pilhofer et al., 2011; Wagstaff & Lowe, 2018); other examples include PhuZ filaments from various bacteriophages (Erb et al., 2014; Kraemer et al., 2012; Zehr et al., 2014). Among these examples, bacterial tubulin bTubA and bTubB show the highest sequence similarity to eukaryotic α*-* and β*-*tubulin (Schlieper et al., 2005), hence the use of the term bacterial microtubules or bMTs. In analogy to eukaryotic MTs, it is proposed that a GTP-cap stabilizes growing bMTs and the filaments exhibit a catastrophe when the GTP-cap is lost (Deng et al., 2017). This dynamic behavior has been reproduced *in vitro* (Deng et al., 2017), although it is currently unknown whether it also occurs *in vivo* and what is its biological role (Diaz-Celis et al., 2017; Kattan et al., 2021). Unlike eukaryotic microtubules, which consist of typically 13 protofilaments, bMTs only consist of 4-5 protofilaments (Deng et al., 2017; Pilhofer et al., 2011) (Figure 1A).

**Figure 1.**
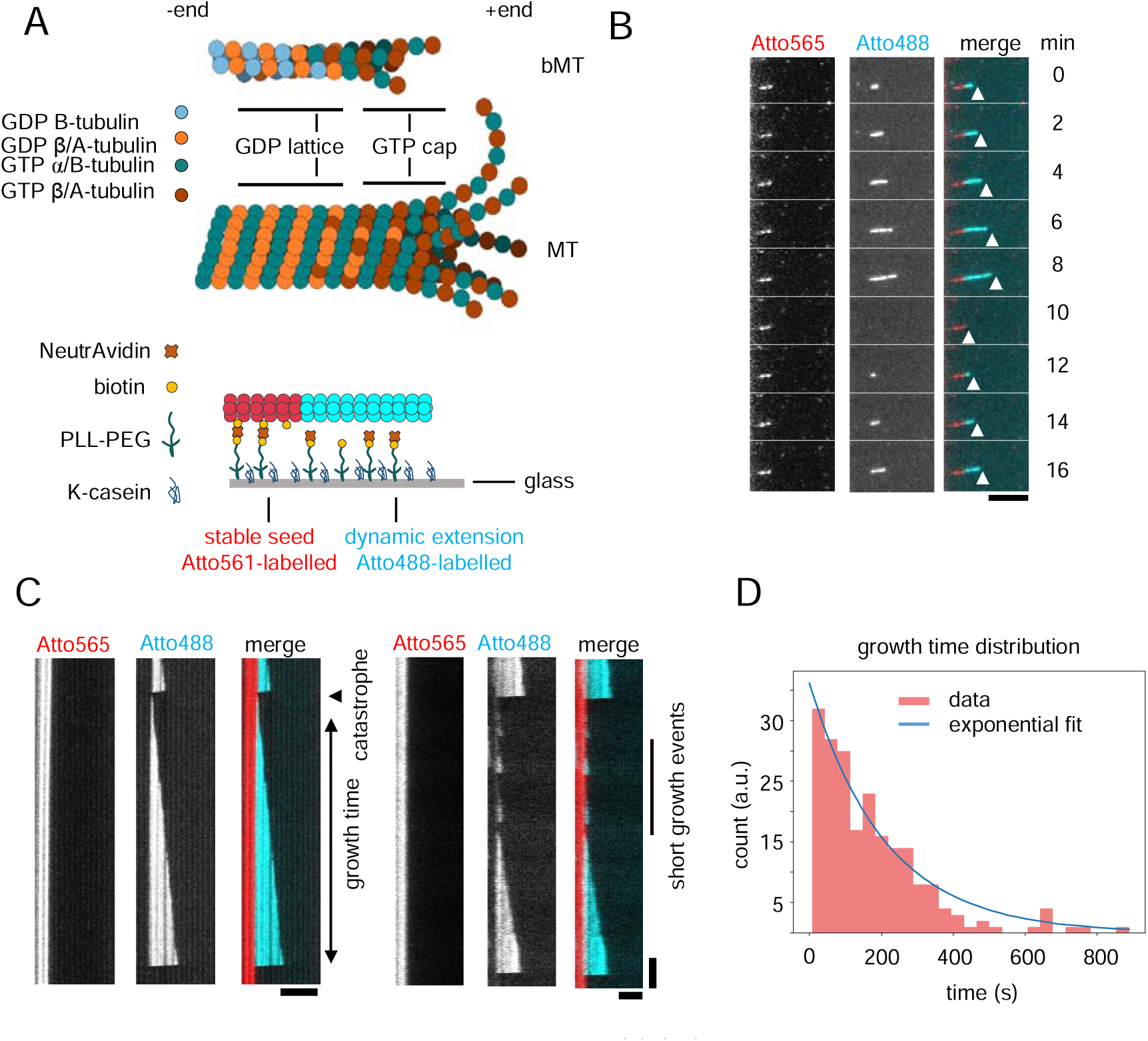
Dynamic properties of free bacterial microtubules. **(A)** (top) Schematic depiction of bacterial and eukaryotic microtubules. Bacterial microtubules are composed of 4-5 protofilaments and both monomers hydrolyze GTP to GDP. Microtubules consist of 13-14 protofilaments instead and only β-tubulin hydrolyses GTP. (bottom) A schematic depiction of the experimental setup. The glass surface is functionalized with PLL-PEG-biotin and NeutrAvidin and passivated with κ-casein. GMPCPP-stabilized, biotin-labeled seeds bind to the surface through NeutrAvidin. Free GTP-tubulin dimers add to the ends of the seeds to generate dynamic filaments. **(B)** A time series of a bacterial microtubule (cyan, Atto488-labeled bTubAB) growing from an immobilized GMPCPP seed (red, Atto565-labeled bTubAB). The tip of the dynamic bMT is marked by an arrowhead over time, showing dynamic instability of the filament. **(C)** Time versus distance graphs (kymographs) of dynamic bMTs (cyan, Atto488-labeled bTubAB) which grow from bMT seeds (red, Atto565-labeled bTubAB). The sudden switch from growth to shrinkage is called catastrophe. Growth time is defined as the growth duration of a filament before a catastrophe. Moreover, on the right, a kymograph example is shown where short growth events were observed to co-exist with longer events. **(D)** A histogram of growth times of free bMTs. The distribution fits the best with an exponential distribution with a mean equal to 209 ± 19 s. Scale bars = 5 µm.

In this paper, we hypothesize that the process that triggers catastrophes in eukaryotic and bacterial MTs may be based on the same basic molecular principles, and that a bMT may ‘simply’ behave as a MT with fewer protofilaments. Studying the dynamic behavior of dynamic bMTs may thus, in addition to its intrinsic interest, provide insight into the dynamic behavior of the structurally more complex eukaryotic MTs. We therefore decided to repeat a series of experiments we recently presented for eukaryotic MTs and ask whether the results on catastrophe statistics can be reconciled using a simple phenomenological model that we have previously introduced for eukaryotic MTs (Kok et al., 2021). In this model, MT growth is presented as a simple one-dimensional process where the size of a stabilizing cap depends on both growth fluctuations of the tip (allowing also for negative growth excursions) and the ra te o f hydrolysis of tubulin-bound GTP incorporated in the lattice. A MT undergoes a catastrophe when the cap is lost due to either or both processes.

We measured the catastrophe statistics both for freely growing bMTs and for force-generating bMTs whose growth was stalled by a microfabricated rigid barrier. In analogy to what was found for eukaryotic MTs, both datasets could be simultaneously fitted using our 1-D model, allowing us to obtain values for the two main fitting parameters of the model: the random GTP hydrolysis rate in the bMT lattice, *K_hyd_*, and the effective diffusion constant describing noisy filament growth, *D_tip_*. Unlike what was previously reported (Deng et al., 2017), we find no evidence for an age-dependency of the bMT catastrophe frequency. In addition, we examined the end-structures of dynamic bMTs using cryo-ET and found tapered ends that show similarity to growing MTs. Interestingly, using cryo-ET we also discovered the existence of bacterial microtubule doublets bMTDs, which may explain the frequent observation of bMT bundles both *in vivo* (Pilhofer et al., 2011) and *in vitro* (Diaz-Celis et al., 2017; Schlieper et al., 2005; Sontag et al., 2005).

## Results

### Dynamic properties of free bacterial microtubules

To assess the dynamics of bacterial microtubules, the surface of a flow chamber was passivated and GMPCPP-stabilized seeds were immobilized using PLL-PEG-biotin and NeutrAvidin. A mixture of 0.88 µM (10:1 unlabeled:labeled) purified bTubAB in the presence of 1 mM GTP was then added to the flow chamber and was imaged at 20°C using total internal fluorescence (TIRF) microscopy (Figure 1A – bottom). We obtained growth dynamics of the filaments by locating the tips of the filaments at 1 s time intervals (Figure 1B and C). Analysis revealed that the average growth velocity and catastrophe frequency of freely growing bMTs were 0.58 *±* 0.02 µm min*^−^*^1^ (mean *±* SEM) and 0.35 ± 0.02 min*^−^*^1^, respectively (Table 1). The catastrophe frequency is calculated by dividing the total number of observed catastrophes by the total observed growth time of the bMTs, where the statistical error is obtained by dividing the frequency by the square root of the number of events (assuming a random catastrophe process). The catastrophe time or growth time, on the other hand, is defined as the time an individual bMT spends growing from nucleation until the filament undergoes a catastrophe (Figure 1C). The growth time distribution of free bMTs was observed to follow an exponential function instead of a gamma function as was reported by (Deng et al., 2017): fitting revealed an exponential time scale of 209 ± 19 s (mean ± SD) (Figure 1D).

**Table 1:** Dynamic instability features and simulation parameters for bMTs and MTs. Dynamic instability parameters: growth speed (mean ± SEM), growth time, and contact time of bMTs and MTs in 0.88 µM bacterial and 15 µM eukaryotic tubulin concentrations, respectively. Growth time and contact time for bMTs is reported as mean ± SEM while it is median ± SE for eukaryotic MTs. Simulation parameters hydrolysis rate (K_hyd_) and diffusion of the tip (D_tip_) (mean ± 95% CI) and N_unstable_. MT data from (Kok et al., 2021).

| | growth speed<br>($\mu\text{m}\cdot\text{min}^{-1}$) | growth time<br>(s) | contact time<br>(s) | $K_{\text{hyd}}$<br>($\text{s}^{-1}$) | $D_{\text{tip}}$<br>($\text{nm}^2\cdot\text{s}^{-1}$) | $N_{\text{unstable}}$ |
| --- | --- | --- | --- | --- | --- | --- |
| bMTs | $0.58 \pm 0.02$ | $173 \pm 10$<br>(n = 218) | $27.9 \pm 1.6$<br>(n = 96) | $0.044 \pm 0.006$ | $500 \pm 151$ | 5 |
| MTs | $1.7 \pm 0.7$ | $155 \pm 15$ | $30.8 \pm 1.3$ | $0.070 \pm 0.006$ | $3720 \pm 310$ | 15 |

### Dynamic properties of stalled bacterial microtubules

We next examined the dynamics of bMTs that were stalled by rigid barriers, using an experimental approach we previously developed for eukaryotic MTs (Kok et al., 2021). The barriers were fabricated on glass coverslips which were then used to make flow chambers. Barriers consisted of a SiO_2_ layer of about 100 nm in thickness, with a SiC overhang to increase the chance for the filaments to interact with the barriers (Figure 2A and B). bMTs which had (near-)perpendicular physical contact with the barriers, were observed to stop growing (stall), buckle, or slide (depicted in Figure 2C – left, middle, and right, respectively). Sliding refers to the situation in which a filament continues to grow while the contact point is sliding along the wall. Buckling, on the other hand, occurs when a filament continues to grow at a fixed barrier contact point. Sliding and buckling were often followed by breaking of the filament. The frequency of observation of each of these events is shown in Figure 2D. As expected and previously shown for eukaryotic MTs, buckling was rare, and stalling was observed mostly for short MTs as it is easier to bend or buckle long MTs (Dogterom & Yurke, 1997).

**Figure 2.**
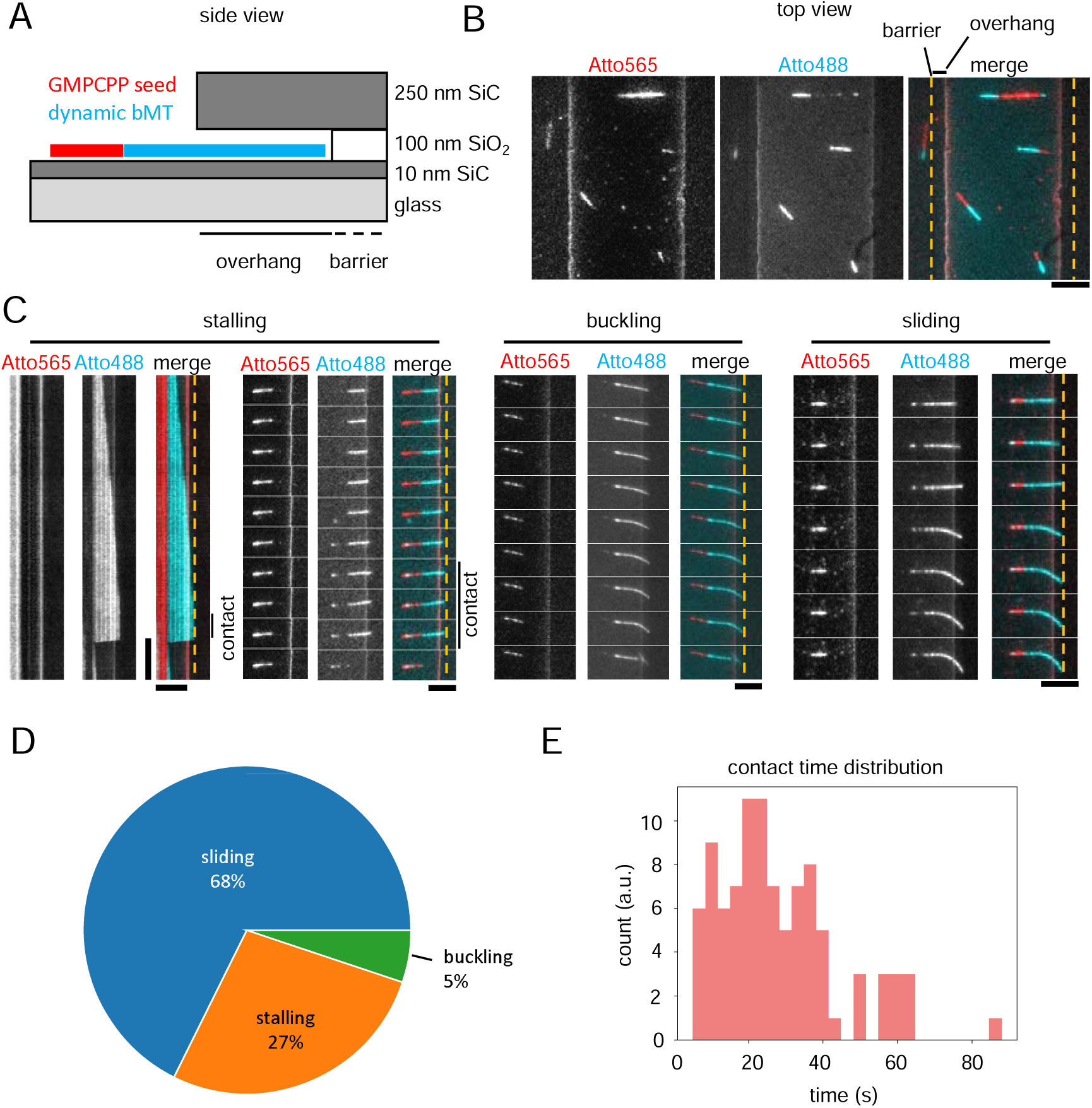
Dynamic properties of stalled bacterial microtubules. **(A)** Schematic illustration of side view of the experimental setup in the presence of micro-barriers. The approximate size of each layer is denoted. In a typical dynamic experiment in the presence of barriers, dynamic bMTs (cyan) grow from immobilized bMT seeds (red) and may interact with a barrier. **(B)** A TIRFM (top) view of an experimental sample, where bMT seeds polymerized in the presence of Atto565-labeled bTubAB are depicted in red, dynamic bMTs polymerized from Atto488-labeled bTubAB in cyan and barriers with yellow dashed lines. **(C)** Various events during a bMT-barrier interaction are shown. (left) A kymograph and a time series of a stalling filament (cyan, Atto488-labeled bTubAB) which grows towards a barrier from an immobilized bMT seed (red, Atto565-labeled bTubAB). (middle) A time series of a buckling bMT which undergoes a breakage eventually. (right) A timeseries of a filament which slides against the barrier and preserves its growth. **(D)** A pie chart of frequencies of the events described in (C) for a total number of 371 events**. (E)** A histogram of stalling or contact times of stalled bacterial microtubules. The average contact time of stalling bacterial microtubules was 27.9± 1.6 s (mean ± SEM). Horizontal scale bars = 5 µm, vertical scale bar = 1 min.

The catastrophe statistics of stalling events were then further analyzed to compare with previous studies done with MTs. As observed for MTs, the situation of net zero growth imposed by contact with the barrier, alters the catastrophe dynamics of the filament. The contact time or stalling time is defined as the time interval during which the net filament growth velocity is equal to zero (v_g_ = 0) while it is in contact with the barrier before a catastrophe occurs. The mean contact time of 96 bMTs was 27.9 ± 1.6 s (mean ± SEM), which is 7.6 times shorter than the average growth time of free bMTs (Table 1). The distribution of contact times for stalled bMTs was not exponential, but clearly peaked as was observed previously for stalled MTs (Figure 2E) (Janson et al., 2003; Kok et al., 2021).

### Catastrophe statistics from simulations based on a one-dimensional model

In analogy to our recent effort for eukaryotic MTs, we then looked for a way to reconcile the measured catastrophe statistics for stalled and free bMTs within a single model. To this end, we simulated microtubule dynamics using a simple one-dimensional MT model that was successfully used before to reconcile the growth time and contact time distributions of free and stalled MTs (Kok et al., 2021).

The model assumes the stochastic hydrolysis of GTP by tubulin subunits within the MT lattice and noisy MT growth, including negative length excursions at the MT end (Figure 3A), which together affect the size of the stabilizing GTP-tubulin cap. The simulation utilizes a Monte-Carlo method and normally requires three fitting parameters: the hydrolysis rate *K_hyd_*, the effective diffusion constant of the tip *D_tip_*, and the minimum number of adjacent GDP subunits at the filament tip needed to trigger a catastrophe *N_unstable_* (Figure 3A).

**Figure 3.**
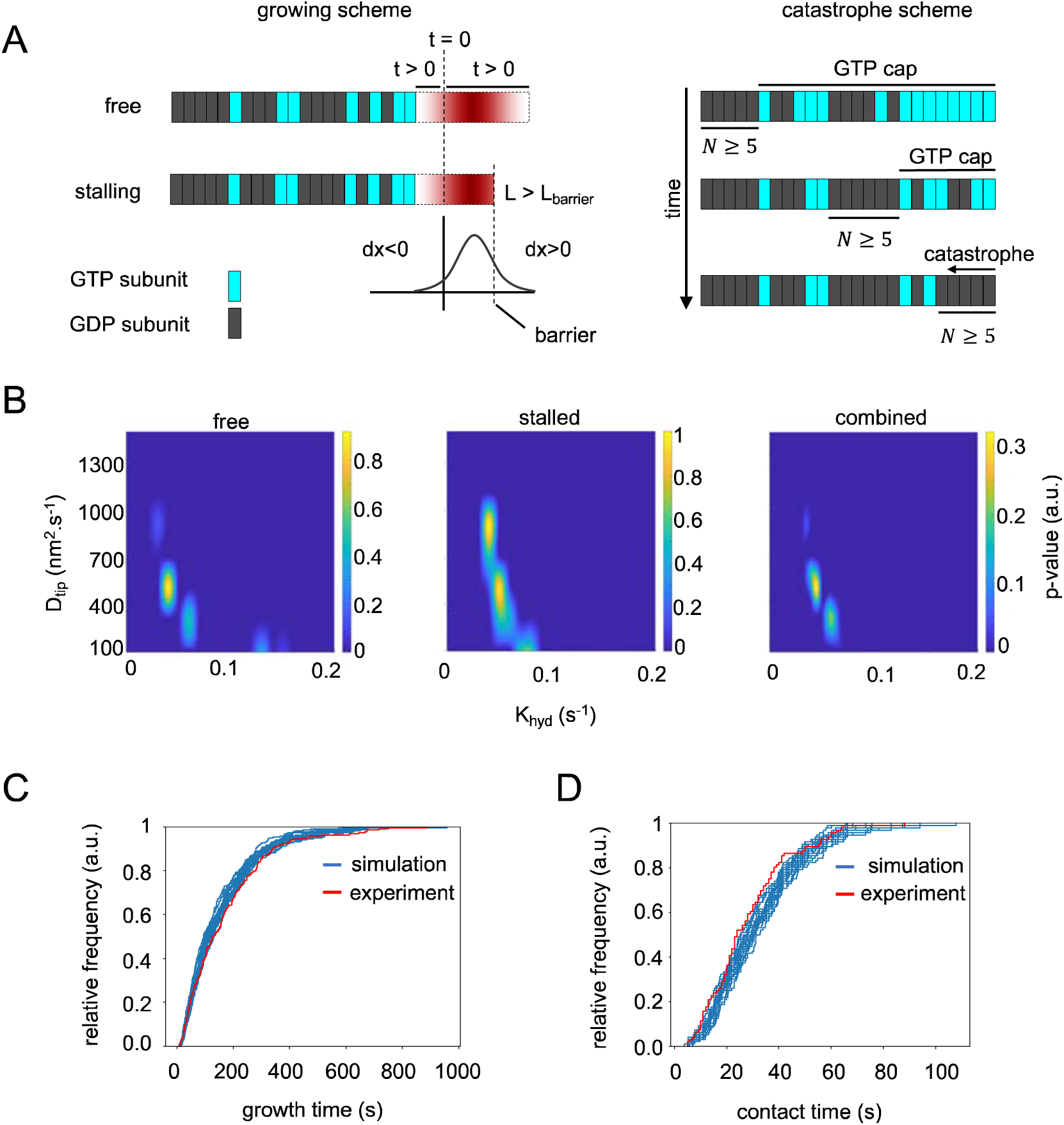
Catastrophe statistics from simulations based on a one-dimensional model. **(A)** A schematic depiction of the basic principles of the one-dimensional model recruited to simulate bMTs. (left) The size and direction of each step in the filament growth at t > 0 is determined by a normal probability density function. During each time step, incorporated GTP-bound subunits (cyan) randomly hydrolyze into GDP-bound subunits (gray) with a rate K_hyd_. Once close to the barrier, the filament length increment is restricted to the distance between the seed and the barrier (L_barrier_). (right) The cartoon represents the process which leads to a catastrophe. A filament undergoes a catastrophe when the number of adjacent GDP-subunits at the end of a filament is equal to or larger than N_unstable_ (N ≥ N_unstable_). **(B)** A heatmap representation of the equality of the simulations and experimental data for free and stalled bMTs and the product of the two (combined). The graphs plot the p-value of a Kolmogrov-Smirnov test by comparing the simulated distribution at different hydrolysis rates and diffusion constants with the experimental data. **(C)** The cumulative distribution of growth times of free bMTs. The experimental data (red) are compared to simulations (blue). 218 bMTs among 1000 simulated filaments were randomly selected and the cumulative distribution is plotted for 25 different selections. **(D)** The cumulative distribution of contact times of stalled bMTs. The red plot and blue plots represent experiments and simulations, respectively. In this case, 25 samples of 96 bMTs among 1000 simulated stalling bMTs were chosen randomly and plotted.

Using fixed parameter values for the growth rate v_g_ = 0.58 µm·min^-1^ (Table 1), subunit length o = 8⁄4 nm (Deng et al., 2017), and maximum filament length given by the measured average distance between the ends of the seeds and the barriers L_barrier_ = 1.45 µm, we sequentially fixed *N_unstable_* at a few different values, i.e., 3, 5, 8, 10, and 12, and then simulated 500 filaments with a range of both *K_hyd_* and *D_tip_*. Distributions of simulated growth times of free bMTs and contact times of stalled bMTs were compared to the experimental data using a Kolmogrov-Smirnov-test (Figure 3B). The fitting parameters which best capture both free and stalled bMTs were found to be *K_hyd_* = 0.044 ± 0.006 s*^−^*^1^ and *D_tip_* = 500 ± 151 nm^2^ s^−1^ (mean ± 95% CI) for N_unstable_ = 5 (Table 1). Figure 3C and D depict the cumulative distributions for the simulated (blue) and experimental (red) datasets. As can be seen from the plots, the simulations closely capture the experimental results.

### Mechanical properties of stalled bacterial microtubules

In addition to analyzing the dynamic properties, we used the stalling data to obtain an estimate of the flexural rigidity of bMTs. When growing filament ends arrive at a barrier, they will continue to grow only when the force needed to buckle the filament is smaller than the force needed to halt the growth process (the stall force) (Dogterom & Yurke, 1997). Since the so-called critical buckling force strongly decreases with filament length, we may assume that the longest observed stalling length corresponds to a situation where the stall force and buckling force are just balancing each other.

For the boundary conditions of our experiment, the buckling force is given by the following equation (Landau et al., 1986):

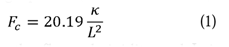

where, *K* is the flexural rigidity and L is the length of the filament. An estimate of the stall force may independently be obtained using the ratio between the subunit concentration during bMT growth and the critical concentration where the subunit on-and off-rates are exactly balanced (Dogterom & Yurke, 1997). Previously, the critical concentration 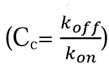 was found to be in the range of 0.4 – 1 µM in bulk measurements (Sontag et al., 2005). Assuming the lower limit (0.4 µM) to be relevant for growth from seeds, and using the value of 0.88 µM for the experimental subunit concentration, the stall force can then be calculated (Dogterom & Yurke, 1997) from the equation:

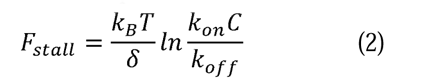

where k_B_ is Boltzmann’s constant, T is the experimental temperature, o is the increase in polymer length per added subunit (2 nm for bMTs), and C is the subunit concentration. Using the above-mentioned values, we obtain a number for F_stall_ on the order of 1.6 pN for bMTs. Using this value for the force in equation **Error! Reference source not found.**, combined with the value for the longest stalling bMT filament in our dataset: L = 5.7 + 0.2 µm, we obtained an approximate value for the flexural rigidity of bMTs: K_bMT_ = 2.6 pNµm^2^. As expected because of their smaller diameter, this value is smaller than that of MTs, see for example the value that was measured for MTs at 23°C and tubulin concentration 20 µM : K_MT_ = 21.2 + 1.7 pNµm^2^ (Janson & Dogterom, 2004).

### Structural properties of bacterial microtubule ends

Dynamic properties of growing MTs have been related to structural properties of their ends. For example, it has been suggested that microtubules in the pre-catastrophe state have extended tapers, stretches of incomplete cylindrical shapes where lateral interaction sites of tubulin subunits are exposed. One hypothesis is that the incomplete set of lateral contacts with neighboring protofilaments destabilizes these long tapers and leads to catastrophe (Coombes et al., 2013; Gudimchuk et al., 2020). Possibly, these tapers are also related to the apparent noise in microtubule growth as incomplete cylindrical structures with reduced lateral contacts may be highly dynamic. Additionally, there is discussion in the literature related to the shapes of the terminal protofilaments that extent from the tapered structures at growing and shortening ends of eukaryotic microtubules: earlier work using single projections of microtubules observed mostly straight protofilaments at growing microtubule ends, and bent protofilaments at shortening ends (Chretien et al., 1995; Mandelkow et al., 1991). However, more recent tomographic analysis challenged this view by showing that growing and shortening microtubule ends carry similarly bent protofilaments, suggesting that GTP-and GDP-tubulin have similar curvatures (McIntosh et al., 2018). To investigate whether the ends of bacterial microtubules resemble eukaryotic microtubules in terms of bent protofilaments and extended tapers, we turned to electron cryo-tomography (cryo-ET) and imaged samples containing bacterial microtubules frozen during growth (Figure 4A). Due to the fast depolymerization of bMTs, we were unable to make samples with filaments frozen in the shortening state.

**Figure 4.**
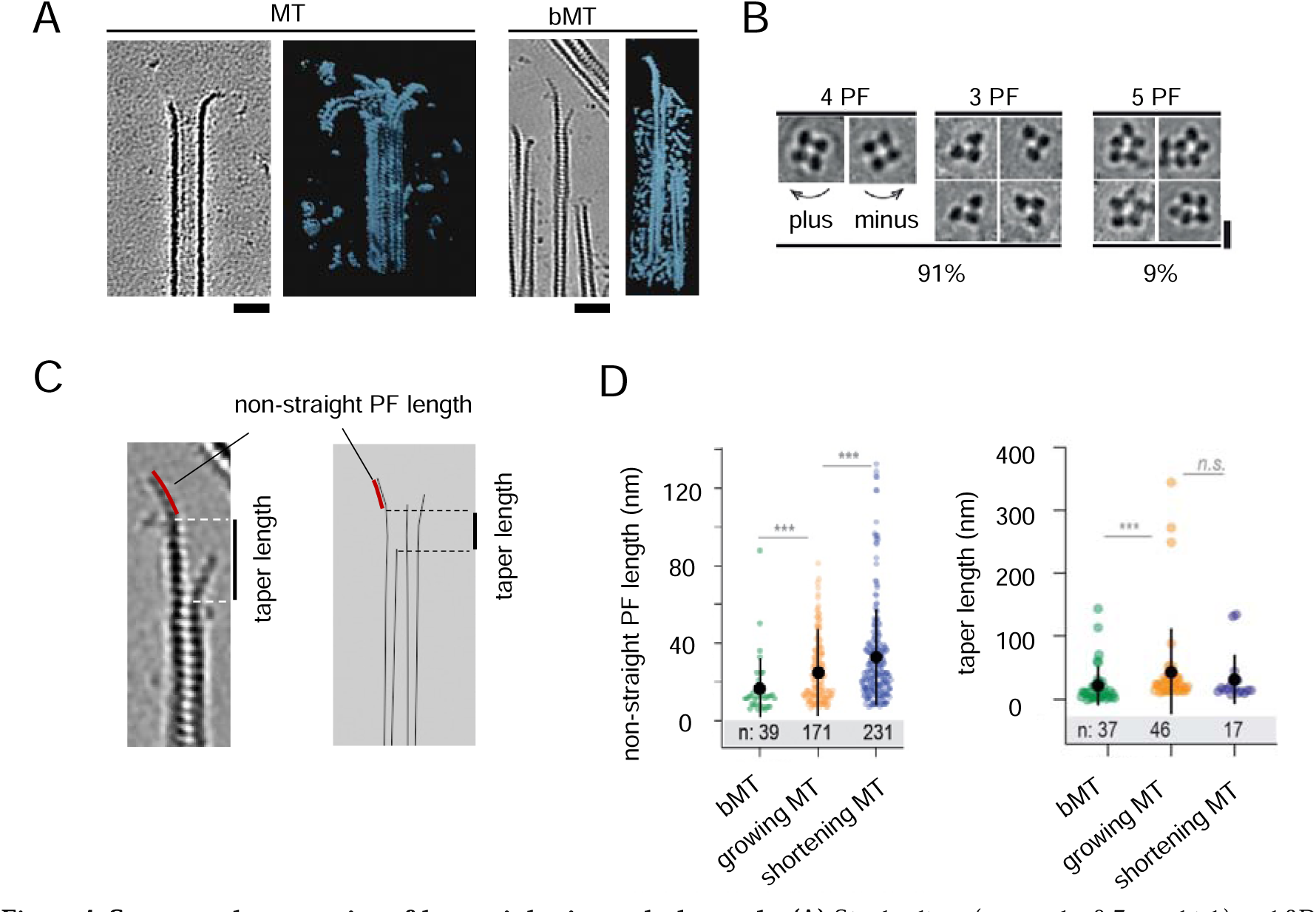
Structural properties of bacterial microtubules ends. **(A)** Single slices (grayscale, 0.7 nm thick) and 3D renders (color) of cryoCARE-denoised tomograms with eukaryotic (left) and bacterial (right) microtubules. Scale bar = 30 nm. **(B)** Axial views of averaged particles used to determine bMT polarity and protofilament number; representative particles for different number of protofilaments and corresponding occurrence of each variant are shown. Due to the asymmetric structure of three-stranded bMTs, we assumed that they were four-stranded which have lost one protofilament and therefore categorized them as four-stranded bMTs. Scale bar = 10 nm. **(C)** A single tomographic slice (left, 0.7 nm thick) and a schematic (right) of a tapered bMT plus end. Taper length and non-blunt protofilament length are defined. **(D)** (left) Length of the bent part of terminal protofilaments of growing bMTs (16.5 ± 2.4 nm (mean ± SEM)) and that of growing and shrinking MTs (24.9 ± 1.2 and 32.6 ± 1.6 nm, respectively (mean ± SEM)). (right) Plus-end taper length of growing bMTs and growing and shrinking MTs. The difference observed between the average taper length of growing eukaryotic and bacterial microtubules was observed to be significant.

To determine the polarity of bMTs, we performed semi-automated particle picking along each filament, and then subtomogram averaging of these particles using dynamo (Castano-Diez et al., 2012) (see Materials and Methods for details). In the axial projections of averaged particles, we observed a mixture of bMTs containing three, four or five protofilaments (Figure 4B). While four-and five-protofilament bMTs appeared to have protofilaments equally spaced around the bMT axis, three-protofilament bMTs looked like four-protofilament bMTs with one protofilament missing, so they were also qualified as four-protofilament ones.

To analyze the shapes of bMT ends, we denoised the reconstructed tomograms using cryoCARE (Buchholz et al., 2019), and then manually segmented the plus-ends to obtain 3D models of the terminal protofilaments (Gudimchuk et al., 2020; Ogunmolu et al., 2022). For comparison, we used plus-ends of eukaryotic microtubules frozen during growth or shortening (see Materials and Methods for details). This analysis revealed that bMTs have tapered ends just like eukaryotic microtubules. However, the majority of bMT plus-ends had protofilaments terminating without a bent part (69%), unlike plus-ends of growing or shortening eukaryotic microtubules that contained 14% and 1% of protofilaments without a bent part, respectively. Focusing on the protofilaments that did terminate with a bent segment, we found that shortening eukaryotic microtubules carried longer bent segments than growing ones, as reported previously (32.6 ± 1.6 nm and 24.9 ± 1.2, respectively; Figure 4C and D) (Gudimchuk et al., 2020). However, bent protofilaments at the plus-ends of bMTs were shorter than observed in both MT samples (16.5 ± 2.4 nm; Figure 4D). Although the differences we found between the lengths of the plus-end tapers in growing: 44.0 ± 10.0 (mean ± SEM) and shrinking: 32.2 ± 9.4 (mean ± SEM) MT samples were not statistically significant, the average taper length of the plus-end of growing bMTs was found to be shorter than that of MTs (22.9 nm ± 5.1; mean ± SEM) (Figure 4D). We conclude that the majority of bacterial microtubules grow with straight ends, unlike eukaryotic microtubules that mostly grow and shrink with flared plus-ends.

### Multi-filament bundle formation of bacterial microtubules

In addition to individual bMT filaments we often observed bundles of various sizes in both cryo-ET and TIRFM images (Figure 5A). TIRFM images suggested that filaments in bundles may be dynamic, as is shown in Figure 5A both in the kymograph and the time series. Cryo-ET data analysis revealed that the filaments in bundles often resembled microtubule doublets, structures that were previously reported for axonemal MTs (Goetz & Anderson, 2010). Bacterial microtubule doublets (bMTDs) were composed of a complete bacterial microtubule of either 4 or 5 protofilaments and an incomplete filament. Different types of doublet structures were observed as represented in Figure 5B. We hypothesize that bMTDs form by the growth of an incomplete bMT alongside a preformed complete bMT. According to our observations, bMT bundles may thus represent different forms of bMTDs as well as bundles of multiple complete bMTs. Note that we preferentially picked thinner bundles for the analysis in Figure 5B. Much thicker bundles are also present in our samples.

**Figure 5.**
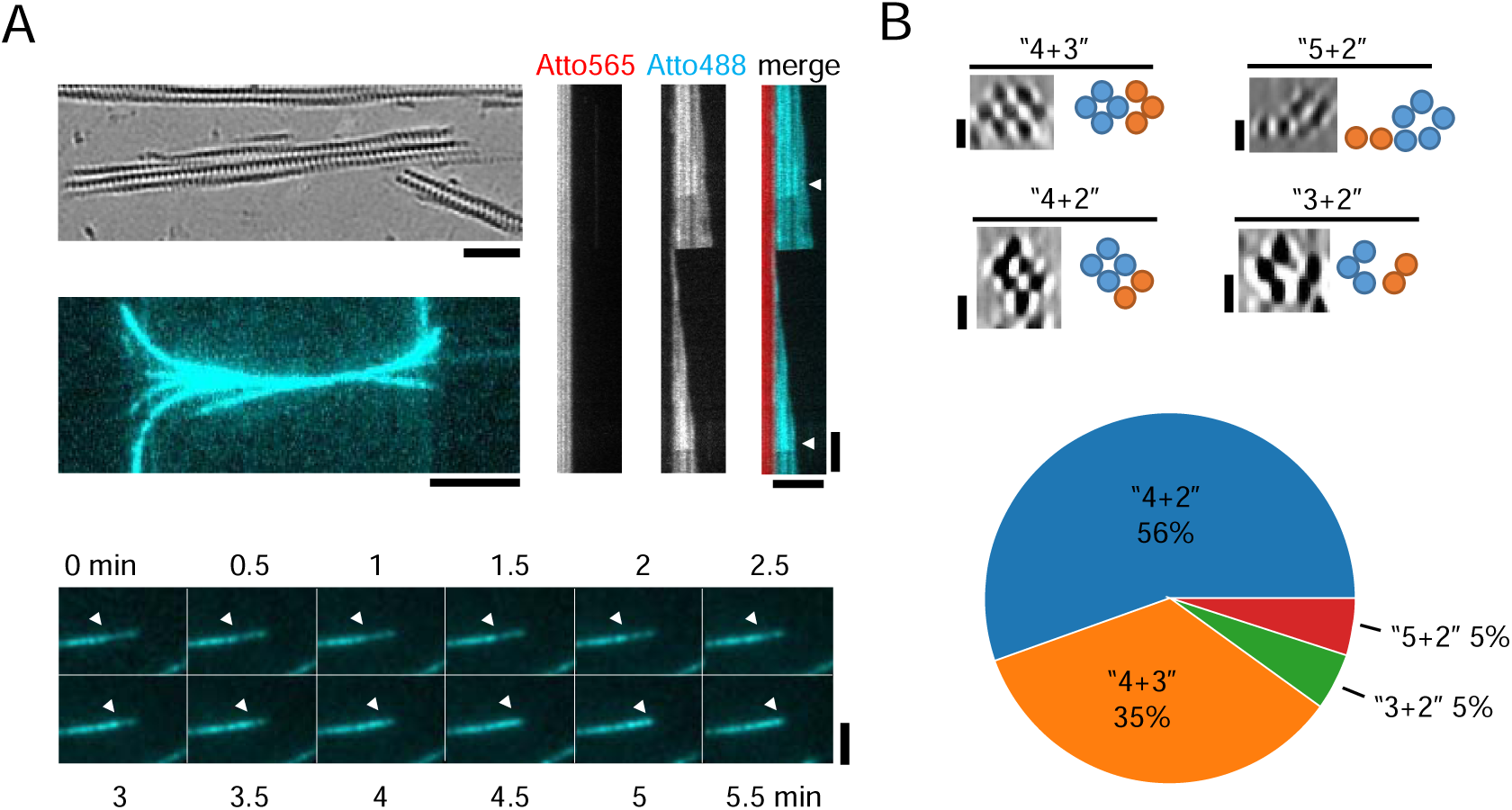
Bacterial microtubules multi-filament bundle formation. **(A)** A single 0.7 nm-thick tomographic slice of a bMT bundle (left-top). Scale bar = 50 nm. A TIRFM image of a bMT bundle growing from bundled seeds in the presence of barriers which cause the filaments bending (left-middle; the seeds are not shown). Scale bar = 5 µm. (bottom) A time series of a growing bMT in a bundle. The arrowheads show the growing tip of the filament in the bundle. Scale bar = 5 µm. (right) A kymograph of a dynamic bundle. The shrinking dynamic filaments are marked with arrowheads. Scale bars: horizontal = 5 µm, vertical = 1 min. The time series image is filtered by Kalman Stack Filter plugin (https://imagej.net/plugins/kalman-stack-filter) in ImageJ for better resolution. **(B)** (top) Axial view of cryoCARE-denoised tomograms showing various conformations of bacterial microtubule doublets and their interpretation. Scale bars = 10 nm. (bottom) A pie chart of the density of bMTD conformations with a sample size of 47 doublets. bMTDs of “5+3” and “4+1” were very rarely observed

## Discussion

In this paper, we studied the dynamic properties of force generating bMTs and compared them with similar experiments performed for eukaryotic MTs, as discussed below. We also used our experiments to obtain approximate values for both the stall force and flexural rigidity of bMTs. The values we obtained, F_stall_ = 1.6 pN and _bMT_ = 2.6 pNµm^2^, are consistent with values obtained for eukaryotic MTs considering that bacterial MTs only consist of 4-5 protofilaments instead of 13. Fewer protofilaments are expected to generate a lower force and thinner tubular structures made of the same material are expected to be less stiff. One may therefore consider bMTs to be similar to eukaryotic MTs in terms of their material properties.

To assess the dynamic properties of bMTs, we analyzed the catastrophe statistics and asked whether they can be reconciled with the catastrophe statistics of eukaryotic MTs within the framework of a simple 1-D model for dynamic instability. Note that in this study we did not vary protein concentration, which is known to affect the dynamic properties of MTs (Gardner et al., 2011; Rickman et al., 2017; Walker et al., 1988) and also bMTs (Deng et al., 2017). Instead, we compared catastrophe data for freely growing bMTs for a specific set of conditions with catastrophe data obtained when growth was completely stalled by a barrier, to be able to make a comparison with the effect that stalling has on the dynamics of eukaryotic MTs. We show that the barrier-bMT interaction strongly enhances the catastrophe frequency of bMTs, as previously found for eukaryotic MTs. In fact, both the average contact time and the distribution of contact times are very similar for stalled bMTs and stalled MTs, as reported in two previous studies (Janson et al., 2003; Kok et al., 2021).

For growing filaments, we expect the average time until catastrophe to be influenced by the growth velocity: faster growing filaments are expected to have a longer growth time (Janson et al., 2003). Therefore, we ideally need to compare bMTs with eukaryotic MTs growing at the same velocity. Indeed, if we compare the growth time we find for growing bMTs to data obtained in one of our older studies (Janson et al., 2003), the results are comparable (although not identical) for similar growth velocities (Figure S1A). It should be noted however that the relation between growth velocity and average growth time (or catastrophe frequency) is known to differ between different eukaryotic tubulin purifications. In our more recent study (Kok et al., 2021), a similar growth time is found at a much higher growth velocity, indicating that these MTs were less stable than the ones in our older study (Figure 1SA).

We next employed a one-dimensional model to reconcile the catastrophe statistics of free and stalled bMTs. Similar to our previous findings for MTs (Kok et al., 2021), we were able to find fitting parameters where the model was able to reconcile both datasets. This suggests that the underlying process that governs the catastrophe statistics of bMTs is similar to that of MTs. To obtain the best possible fits, we had to assume that the minimum number of adjacent GDP-bound subunits at the filament tip necessary for a catastrophe N_unstable_ = 5, which corresponds to one layer of bTubAB dimers. This is in accordance with what was previously reported for MTs (N_unstable_ = 15, corresponding to about one layer of tubulin dimers).

The first main fitting parameter, *D_tip_*, which represents the diffusive or noisy growth of the filament, was found to be *D_tip_*= 500 ± 151 nm^2^ s*^−^*^1^. Interestingly, this is of similar order of magnitude as the diffusion constant obtained from a model fit for eukaryotic MTs growing at a velocity comparable to that of our bMTs and also to numbers experimentally obtained by (Gardner et al., 2011) (Figure S1B). However, for faster growing MTs in our recent study we find a much higher value for this parameter (*D_tip_* = 3720 ± 310 s^-1^), corresponding to a much shorter average experimental growth time (Kok et al., 2021). We note that catastrophe frequencies (and therefore average experimental growth times) vary widely between different tubulin batches which may thus be related to differences in diffusive growth behavior (possibly due to structural differences) at MT ends.

Interestingly, while we can reconcile the different growth times of freely growing MTs for our different datasets only by assuming different values for *D_tip_*, it appears that the catastrophe statistics of stalled filaments is similar for all datasets (bMTs as well as MTs for both our previous studies). As can be seen in Figure 3B (middle panel), the quality of the fit for stalled filaments is not very sensitive to the value of *D_tip_*, but instead largely determined by the hydrolysis rate *K_hyd_* (see also Figure 4A, middle panel in (Kok et al., 2021)). This parameter was found to be of the same order of magnitude for both bMTs and MTs, consistent with similar catastrophe statistics for stalled filaments. The obtained value for bMTs, *K_hyd_* = 0.044 ± 0.006 s*^−^*^1^, is higher than the bulk hydrolysis rate previously measured by Diaz-Celis et al.: *K_hyd_* ≈ 0.02 GTP. s*^−^*^1^. [bTubAB]^-1^ (Diaz-Celis et al., 2017; Sontag et al., 2005). This difference may be expected since the polymerization of bacterial tubulin is likely to catalyze the hydrolysis of GTP. The GTP hydrolysis rate for eukaryotic MTs has also been measured and was found to be at least: 0.04 s^-1^ (O’Brien et al., 1987).

We find that the growth times of free bMTs are exponentially distributed, which is contradictory to previous findings which suggested a lack of short events in the growth time distribution of free bMTs and hence a gamma-distributed growth time (Deng et al., 2017). As can be seen from a sample kymograph of a bMT (Figure 1C) there is a considerable number of short (less than three pixels in length) filaments (cyan) growing from stabilized seeds (red). Although the length of these filaments was at the order of the resolution of the microscope (250 nm) their lifetime was long enough to be measured. A gamma fit revealed a gamma shape parameter equal to 1.4 ± 0.1. However, the lowest Bayesian information criterion (BIC) for fitting a gamma distribution to the observed growth times was 167 (Figure S2A) which is higher than the lowest BIC for the exponential fit: 124 (Figure 1D). We also found the taper length of bMT filaments to be independent of length and hence age (Figure S2B). Nevertheless, a slight negative correlation with a Spearman’s correlation coefficient of −0.14 is noticeable which could be a result of the low number of data points. Note however that the length range that could be measured for bMT filaments was limited: a high density of filaments and a limited field of view in our experiments make it difficult to track the full length of a filament.

Finally, bacterial microtubules are highly prone to form bundles (Figure 5). We find that complete and incomplete bMTs may coexist in bacterial microtubule bundles. The combination of a complete and one or more incomplete bMTs forms bacterial microtubule doublets (bMTDs). bMTDs were observed and reported previously. However, these bundles were called “protofilament bundles” disregarding the presence of complete microtubules (Schlieper et al., 2005; Sontag et al., 2005). How these structures are formed and whether each of these sub-structures is dynamic is not known. Schmidt-Cernohorska et al. have recently shown that by removing the C-terminal tail residues of [z]β-tubulin, a MT doublet could form (Schmidt-Cernohorska et al., 2019). We may thus speculate that the main reason for bundling and formation of bMTDs is the absence of highly acidic C-terminal tails in bTubA and B (Martin-Galiano et al., 2011; Schmidt-Cernohorska et al., 2019).

## Acknowledgments

We thank Anne Doerr for protein purification and labeling, Maurits Kok for providing the Matlab scripts, and Florian Huber for his assistance during working on simulations. We would like to thank Wiel Evers (TUDelft), and Christopher Diebolder and Wen Yang (NeCEN) for their excellent technical assistance in data acquisition for cryo-ET. The cryo-ET data on eukaryotic microtubules were collected at the Netherlands Centre for Electron Nanoscopy (NeCEN) made possible through financial support from the Dutch Roadmap Grant NEMI (NWO.GWI.184.034.014). We also acknowledge Tomohiro Shima and Hanjin Liu for the discussions on cryo-ET experiments. This work was supported by The Netherlands Organization of Scientific Research (NWO/OCW) Gravitation program Building a Synthetic Cell (BaSyC) (024.003.019). V.A.V. acknowledges support from the QMUL Startup grant SBC2VOL8.

## Author Contributions

R.A.H fabricated micro-fabricated barriers, performed experiments, analyzed the data, and ran the simulations. V.A.V. performed and analyzed cryo-ET experiments. All authors wrote the paper. M.D. coordinated the project.

## Declaration of interests

The authors declare no competing interests

## STAR Methods

### Protein purification

*Prosthecobacter dejongeii* bacterial tubulins A and B were co-expressed and purified following the procedure explained by (Diaz-Celis et al., 2017) except that vectors were overexpressed in OverExpress C41(DE3) Chemical Component Cells (Immunosource) instead of BL21(DE3). In short, following the steps of harvesting the cells, lysing, and removing the debris, the proteins went through three consecutive polymerization-depolymerization cycles using Beckman ultracentrifuges. Each cycle started with addition of 5 mM MgCl_2_ and 0.5 M KCl and incubation for 10 min at 25°C. 2 mM GTP was then added to the solution and the solution was immediately centrifuged at 100 000×g for 30 min in a Type 60 Ti Beckman rotor (273 000 rpm). The supernatant was discarded, and the pellet was resuspended with 3 ml ice-cold 50 mM HEPES-KOH pH 7 and incubated for 30 min on ice with occasional pipetting. A cycle was completed by a 100 000×g centrifuge for 30 min at 4°C in the similar rotor. The pellet was discarded, and the supernatant was warmed to 25°C for the next cycle.

The pellet of the final cycle was resuspended in 500 µl (instead of 3 ml) 50 mM HEPES-KOH pH 7, incubated for 30 min on ice with occasional pipetting. Final concentration was measured by a Nanodrop^TM^ 2000 spectrophotometer at 280 nm (A280). Extinction coefficient = 103754.2 M^-1^cm^-1^. In the end, aliquots of 1 µl were made and snap frozen in liquid nitrogen and stored in −80°C for later use.

### Protein labeling

Purified bTubAB were labeled on their lysine residues either by biotin or Atto dyes following the procedure described by (Deng et al., 2017). Inactive proteins were first discarded from the bTubAB solution by completing a polymerization-depolymerization cycle started by adding GTP and MgCl_2_ to a bTubAB solution at final concentrations 2 mM and 5 mM, respectively, and incubating for 10 min at 37°C. The solution was centrifuged in a Beckman airfuge rotor at ambient temperature at 30 psi for 5 min. The supernatant was discarded, and the pellet was resuspended in buffer L (80 mM PIPES-KOH, 75 mM K-Acetate, 0.5 mM EDTA, and 0.2 mM TCEP pH 7.5). The label (in DMSO) at a ratio of 1:2 (protein:label) in the presence of 2 mM GTP was added to the solution and incubated for 30 min at room temperature. Then the mixture was supplemented by 75 mM K-Glutamate and 2 mM GTP and was gently added as a layer on top of a pre-warmed glycerol cushion (3 ml buffer L, 1.2 ml glycerol, 225 µl K-Glutamate (2 M) and 1.575 ml MilliQ) and centrifuged at 100 000×g in a Type 60 Ti Beckman rotor (27 300 rpm) at 35 °C for 30 min. After removing the cushion and washing the pellet with warm MRB80 (80 mM PIPES pH 6.8, 4 mM MgCl_2_, 1 mM EGTA), the pellet was re suspended in 50 µl MRB80 on ice for 10 min and centrifuged in a Beckman airfuge on cold rotor for 5 min at 30 psi. At the end, we collected the supernatant and measured the concentration as well as the labeling ratio with a Nanodrop^TM^ 2000 spectrophotometer using A280 and then A500 to correct the label absorption at 280 nm. The aliquots were then snap frozen in liquid Nitrogen and stored at *−*80 °C.

### Fabrication of micro-barriers

Micro-barriers were fabricated by adapting the procedure introduced by (Kok et al., 2021). The procedure started by cleaning 24×24 mm coverslips in a base piranha solution containing H_2_O_2_:NH_4_OH:H_2_O with 1:1:5 ratio at 75 °C. Using a PE-CVD machine, three layers were coated on each coverslip; first a SiC layer of about 10 nm, then a SiO_2_ of about 100 nm and on top of all, a roughly 250 nm layer of SiC. On the next step, we transferred the barrier pattern using lithography with positive photoresist S1813 and exposed with near-UV light source for 5 s. The photoresist was developed by MF321 for 1 min.

Afterwards, the photoresist-free surfaces were etched by an RIE plasma etcher (Sentech Etchlab 200) with a gas mixture of 50 sccm:2.5 sccm for CHF_3_:O_2_ for 1180 s. This resulted in the removal of the top SiC layer entirely and the SiO_2_ layer partially. The chamber pressure and the RF generator power were set to 50 µbar and 50 W, respectively; the bias voltage was 200 V, as a result. Photoresist residues were removed either by wet or dry cleaning afterwards. The wet cleaning was done by dipping into Acetone for 10 min, and for dry cleaning was done by an oxygen plasma cleaner at 600 W and oxygen pressure of 600 sccm for 1 h. We employed dry cleaning instead of wet cleaning because wet cleaning was not always reliable. However, unexpected wrinkles could form at the overhangs possibly because of the heat produced by the plasma cleaning device. Nevertheless, wrinkles were not present in any of the samples used in this study. At the end, by etching through the exposed SiO_2_ layer by dipping into buffered hydrofluoric acid (HF:NH_4_F with ratio 12.5:87.5%) for 5 min an overhang of about 1.5 µm length was created.

### Reconstitution of bacterial microtubule dynamics

Bacterial microtubule dynamic assays were inspired by (Deng et al., 2017) and eukaryotic microtubule dynamic assays in (Kok et al., 2021). As mentioned in the “protein purification” section, bTubAB buffer was 50 mM HEPES-KOH pH 7 while rest of the ingredients were dissolved and stored in MRB80 (80 mM PIPES pH 6.8, 4 mM MgCl_2_, 1 mM EGTA). When necessary, the protein dilution was done in MRB80 as well.

#### bMT seeds

With two main considerations in mind, i.e., minimizing the bundling effect (Diaz-Celis etal ., 2017) and maximizing the ultimate seed length, 3 µM (67% unlabeled, 7% Atto-565-labeled, and 26% biotin-labeled) bTubAB in the presence of 10 mM KCl in MRB80 was centrifuged in an airfuge (Beckman) cold rotor at 30 psi for 5 min, the pellet was discarded, and the supernatant was polymerized in the presence of 0.5 mM GMPCPP for 10 min at 37 °C. Seeds were made specifically for each experiment.

#### Flow chambers

Each coverslip with barriers was cleaned with oxygen plasma (50 W, 200 mbar) for 5 min. Then, using double-sided tape straps, a 10 µl flow chamber was made on a glass slide with barriers being perpendicular to the flow direction. The chamber was functionalized in three consecutive steps with 0.2 mg·ml*^−^*^1^ PLL-PEG-biotin (20 %), 0.2 mg·ml*^−^*^1^ NeutrAvidin and 0.5 mg·ml*^−^*^1^ κ-casein with an incubation time of 10 min at room temperature per step. The channel was rinsed with 20 μl of MRB80 after each incubation and at the end with 20 μl of 0.2 % methyl cellulose in MRB80. Afterwards, 20 μl of seed mixture was added to the chamber and incubated for 10 min at room temperature in dark. The incubation time was chosen long enough for an even distribution of seeds. Not adhered seeds were washed away twice with 0.4 µM bTubAB supplemented with 2 mM GTP and 0.2 % methyl cellulose to prevent seed breakage and fast depolymerization.

#### Dynamic filaments

An oxygen-scavenging system composed of 4 mM DTT, 200 µg ml*^−^*^1^ catalase, 400 µg ml*^−^*^1^ glucose oxidase, and 50 mM glucose was used in the fluorescent imaging to minimize photobleaching effects. 20 µl of tubulin mixture composed of 0.88 µM bTubAB (including 0.08 µM Atto-488-bTubAB), the oxygen scavenging system, 0.5 mg·ml*^−^*^1^ κ-casein, 1 mM GTP, and 0.2 % methyl cellulose was centrifuged in an airfuge (Beckman) on an ice-cold rotor at 30 psi for 5 min, warmed in hand and gently added to the chamber. The chamber was sealed with vacuum grease and imaged immediately at 20°C.

### Imaging and image treatment

All the images were acquired using an inverted Nikon Eclipse Ti-E microscope with a perfect focus system. The Nikon Plan Apo λ 100x NA 1.45 oil immersion objective was equipped with an extra lens which magnified the images to a pixel size of 107 nm. The total internal reflection fluorescent (TIRF) system was equipped with a FRAP/TIRF Ilas2 system (Gataca Systems), a 150 mW 488 nm, and a 100 mW 561 nm laser. Image acquisitions were performed at 1 Hz for 20 min with an exposure time of 100 ms and were controlled with MetaMorph 7.8.8.0 software (Molecular Devices). All the image acquisitions were performed within 90 min from the GTP tubulin being introduced to the chamber.

The drift assigned to the image stacks during image acquisition was removed by subpixel image registration through cross correlation. Corrected images were then used to measure the dynamics of bMTs utilizing Fiji (https://imagej.net) software. All kymographs were created by KymoResliceWide plugin (https://imagej.net/plugins/kymoreslicewide) with a straight line of 9 pixels (0.95 µm) width drawn along the long axis of bMTs. The position of the tip of the filaments in the kymographs were used to measure the growth rate by employing a Matlab script. In the process of analyzing the TIRF images, bundles were discarded from the dataset manually.

### Simulations

The simulations were performed following the procedure introduced by (Kok et al., 2021). To execute the simulations, we first embedded fixed parameters: growth speed V = 0.58 µm/min, subunit size 0 = 2 nm, and average distance of stalling bMTs from barriers L_barrier_ = 1.45 µm to the simulations. To measure L_barrier_, bundled filaments were manually discarded from dataset.

Fitting parameters were then identified by comparing the growth times and contact times of simulated bMTs to the experimental data employing a Kolmogrov-Smirnov-test (KS-test) using a Matlab (R2019) code. The test quantifies the equality of empirical cumulative distribution function of the two samples. The equality is then determined by a number (p-value) between 0 and 1 where a higher p-value represents a better fit. We first fixed the N_unstable_ at a few values (i.e., 3, 5, 8, 10, and 12) and ran the simulations for 500 bMTs at fixed rage of K_hyd_ and D_tip_. The ranges for hydrolysis rate (K_hyd_) and diffusion constant of the tip (D_tip_) were inspired by known values for both bMTs and MTs and cubic interpolation was applied to those values. The fitting parameters which simultaneously satisfy free and stalled bMTs’ behavior refer to the highest value of the product of the p-values of the two situations (Figure 3B). The parameter “max_MT_growth_time” which refers to the number of time steps in the simulations was set to 5300.

After exploring the best fitting parameters (Table 1), these parameters were recruited to simulate 1000 bMTs for each experimental situation i.e., freely growing and stalling. To visualize the similarity between the results of simulations and experiments, we randomly sampled bMTs and plotted the empirical cumulative distribution for each sample (Figure 3C and D). Each sample had the same size as the experimental sample size for free and stalled bMTs i.e., 218 and 96, respectively. The open-license simulations code is available on GitHub (https://github.com/florian-huber/mtdynamics).

### Sample preparation for cryo-ET

Bacterial microtubules were polymerized in 80 mM K-PIPES (pH 6.8, supplemented with 25 mM KCl, 1 mM EGTA, 4 mM MgCl_2_ and 2 mM GTP). 20 µM bTubAB were polymerized for 10 min at 23°C and then for about 5 min in presence of 5 nm gold beads. 3.5 µL of this solution was added onto a freshly glow-discharged lacey carbon grid suspended in a chamber of a Leica EM GP2 plunger equilibrated at 98% relative humidity and 26°C, and immediately blotted from the back side for 4s, and plunge-frozen in liquid ethane. The grids were stored in LN_2_ until further use.

Growing eukaryotic microtubules were prepared by polymerizing 15 µM tubulin in presence of freshly prepared GMPCPP-stabilized microtubule seeds for 15 min in a buffer containing 80 mM K-PIPES pH 6.9, supplemented with 80 mM KCl, 1 mM EGTA, 4 mM MgCl_2_ and 1 mM GTP. Grids were prepared as described above.

Shortening eukaryotic microtubules were prepared by first attaching DIG-labelled, GMPCPP-stabilized microtubule seeds to freshly glow-discharged, holey carbon grids pre-incubated with anti-DIG IgG. After 1 min incubation with the seeds, a grid was suspended in the chamber of a Leica EM GP2 plunger equilibrated at 98% relative humidity and 35°C. Microtubules were polymerized on the grid following the addition of 3 µL of 20 µM tubulin in a buffer containing 80 mM K-PIPES pH 6.9, 1 mM EGTA, 4 mM MgCl_2_ and 1 mM GTP. After 10 min of growth, the shortening was initiated by diluting the tubulin on the grid using 30 µL of the same buffer with 5 nm gold beads but without tubulin, prewarmed at 37°C. The grids were blotted and plunge-frozen in liquid ethane 45s after the dilution to allow all microtubules to switch to shortening.

### Recording, reconstruction and denoising of tomograms

Bacterial microtubules were imaged using a JEM 3200FSC microscope (JEOL) equipped with a Gatan K2 Summit electron detector and an in-column energy filter operated in zero-loss imaging mode with a 30 eV slit width. Images were recorded at 300 kV with a nominal magnification of 10000x, resulting in the pixel size of 3.668 Å at the specimen level. Automated image acquisition was performed using SerialEM 3.8.5. software (Mastronarde, 2005). Two samples of growing microtubules were imaged in the same way, while one sample was imaged using a Titan Krios microscope equipped with a Gatan K3 electron detector at the Netherlands Center for Electron Nanoscopy (NeCEN, Leiden, Netherlands).

Images were recorded at 300 kV with a nominal magnification of 26000x, resulting in the pixel size of 3.28 Å at the specimen level. Automated image acquisition was performed using SerialEM. Shortening eukaryotic microtubules were imaged using a Titan Krios microscope equipped with a Gatan K2 electron detector (NeCEN). Energy filtering was performed at post-processing. Automated image acquisition was performed using Tomography software (Thermo Fisher). Images were recorded at 300 kV with a nominal magnification of 33000x, resulting in the pixel size of 4.24 Å at the specimen level. In all cases, we recorded bidirectional tilt series ranging from 0° to ±60° with tilt increment of 2°; a total dose of 100 e−/Å2 and the target defocus set to -4 µm.

Tomograms were reconstructed and denoised as described previously (Maan et al., 2022). In brief, motion was corrected using MotionCor2 (Zheng et al., 2017), tomographic volumes were reconstructed using IMOD 4.11 (Kremer et al., 1996) with twofold binning. Denoising was performed using cryoCARE (Buchholz et al., 2019) on tomographic volumes reconstructed using identical paremeters from odd and even frames using python scripts (available at https://github.com/NemoAndrea/cryoCARE-hpc04).

### Analysis of tomograms

To determine polarity of bacterial microtubules, subvolumes containing filaments of sufficient length were manually extracted from non-denoised tomographic volumes using 3dmod, and averaged using dynamo (Castano-Diez et al., 2012). To create 3D averages, particles were cropped every 4 nm using a “Filament With Torsion” model, and subsequently averaged. Cross-sections of bMTs presented in Fig 5C show sum projections of particles averaged from a single filament. Based on this analysis, we were able to successfully determine polarity of 70 out of 89 filaments; filaments with unclear polarity were discarded from further analysis. Polarity of eukaryotic microtubules was determined by analysing their moiré patterns (Chretien et al., 1996).

To determine protofilament lengths, denoised subvolumes containing plus-ends were manually segmented as described (Gudimchuk et al., 2020; Ogunmolu et al., 2022). Coordinates of tubulin subunits in a protofilament were manually placed on a 3D model in 3dmod by inspecting a subvolume opened in *slicer* and *isosurface* windows of 3dmod side by side, to monitor the accuracy of manual segmentation on a 3D rendered volume. Protofilament lengths were then extracted from manually segmented models using *howflared* and analyzed using MATLAB scripts (https://github.com/ngudimchuk/Process-PFs). Taper length was determined as the distance between the first bent segments of the protofilaments bending closest to and farthest from the end of a filament.

## Supplemental Information titles and legends

**Figure S1: Growth times and diffusion constants for bacterial and eukaryotic MTs. (A)** Average experimental growth times versus average growth speeds for bMTs and MTs from various studies. The graph indicates that the average growth time of bMTs (red) is comparable to that of MTs for similar growth speed in our older study (Janson et al., 2003) (blue). However, the growth time of MTs in our more recent study (Kok et al., 2021) (magenta) shows a much lower growth time at a higher growth speed. **(B)** Diffusion constant versus growth speed for bMTs and MTs from different studies. The diffusion constant for bMTs obtained from simulations in this study (red) is comparable to both measured (Gardner et al., 2011) (green) and simulated (Janson et al., 2003) (blue) diffusion constants for MTs at similar growth speed.

**Figure S2: Bacterial microtubule lifetime is age independent. (A)** A gamma distribution function is fitted to the observed growth times of free bMTs. The shape parameter of the gamma distribution was 1.4 ± 0.1. **(B)** Tapering length of bMTs versus the filament’s length. The sample included 20 µM GTP-bTubAB and was imaged by the cryo-ET technique. Blue circles refer to the filaments of full length and orange circles represent the bMTs which were longer than the field of view and the length refers to the length of the visible segment. The average tapering length was 22.9 ± 5.1 nm. A slight negative correlation (Spearman correlation coefficient = − 0.14) is noticeable.

## Notes

### Competing Interest Statement

The authors have declared no competing interest.

