## Supplemental figures for "Dynamic instability of force-generating bacterial microtubules"

### Slide 1
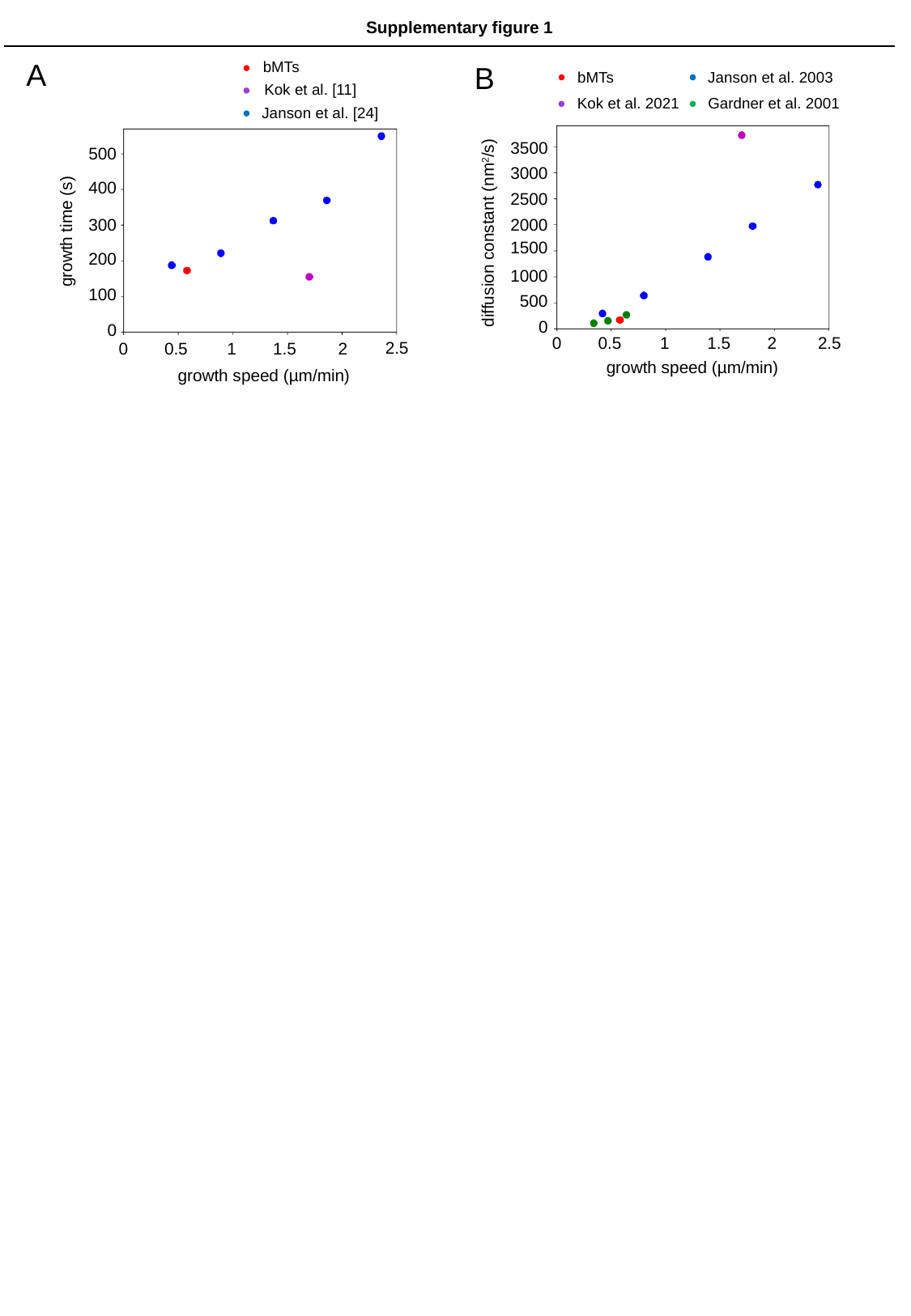

Supplementary figure 1
A
bMTs
Kok et al. [11]
Janson et al. [24]
500
400
300
growth time (s)
200
100
0
2.5
0
0.5
1
1.5
2
growth speed (µm/min)
B
Janson et al. 2003
bMTs
Gardner et al. 2001
Kok et al. 2021
3500
3000
2500
2000
diffusion constant (nm^2^/s)
1500
1000
500
0
0
0.5
1
1.5
2
2.5
growth speed (µm/min)

### Slide 2
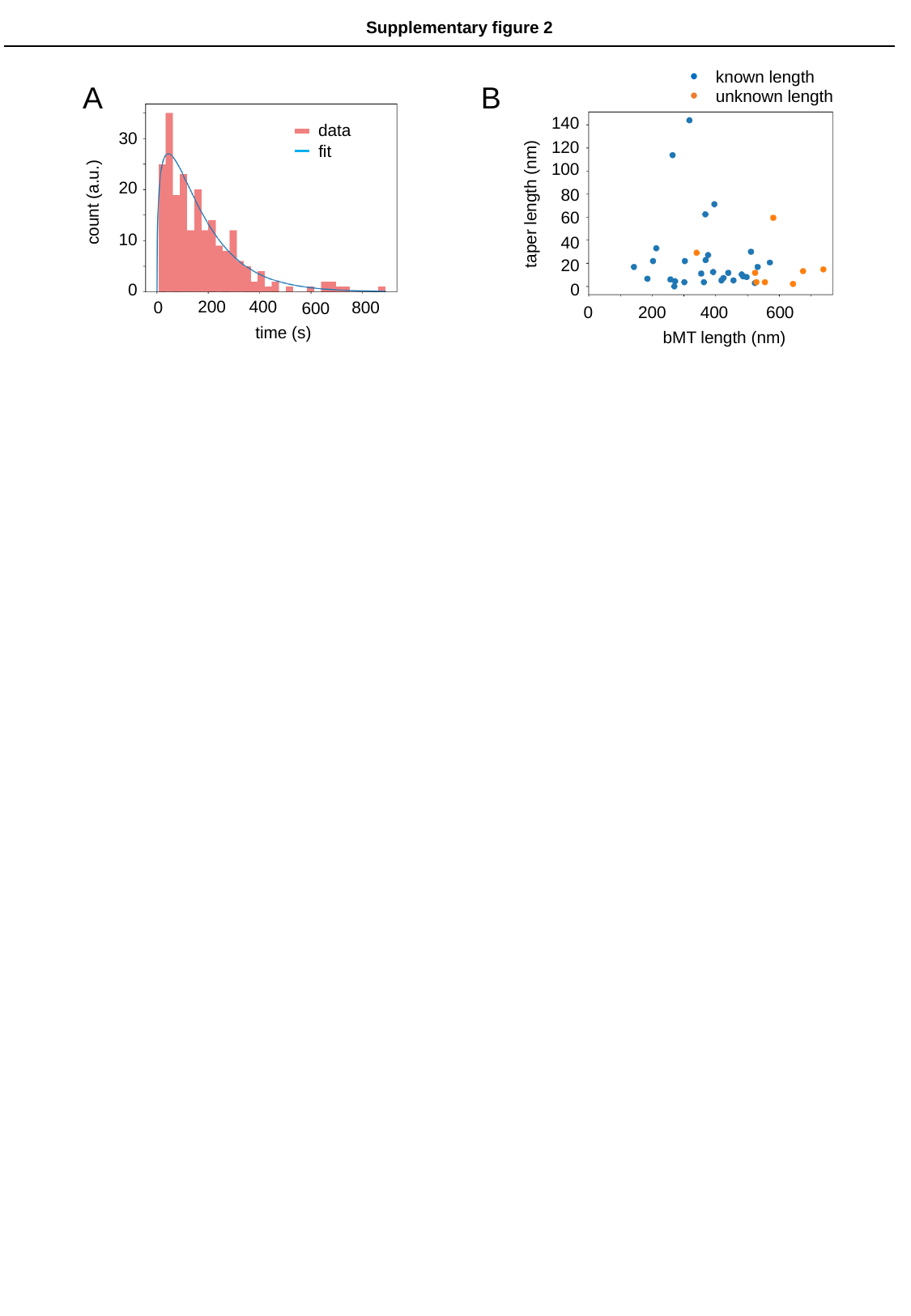

Supplementary figure 2
known length
unknown length
taper length (nm)
bMT length (nm)
140
120
100
80
60
40
20
0
0
200
400
600
B
A
data
30
fit
20
count (a.u.)
10
0
200
400
0
800
600
time (s)
